# Diffraction contrast in cryo-scanning transmission electron tomography reveals the boundary of hemozoin crystals *in situ*

**DOI:** 10.1101/2022.05.13.491750

**Authors:** Debakshi Mullick, Katya Rechav, Leslie Leiserowitz, Neta Regev-Rudzki, Ron Dzikowski, Michael Elbaum

## Abstract

Malaria is a potentially fatal infectious disease caused by the obligate intracellular parasite *Plasmodium falciparum*. The parasite infects human red blood cells (RBC) and derives nutrition by catabolism of hemoglobin. As amino acids are assimilated from the protein component, the toxic heme is released. Molecular heme is detoxified by rapid sequestration to physiologically insoluble hemozoin crystals within the parasite’s digestive vacuole (DV). Common antimalarial drugs interfere with this crystallization process, leaving the parasites vulnerable to the by-product of their own metabolism. A fundamental debate with important implications on drug mechanism regards the chemical environment of crystallization *in vivo*, whether aqueous or lipid. This issue had been addressed previously by cryogenic soft X-ray tomography. We employ cryo-scanning transmission electron tomography (CSTET) to probe parasite cells throughout the life cycle in a fully hydrated, vitrified state at higher resolution. During the acquisition of CSTET data, Bragg diffraction from the hemozoin provides a uniquely clear view of the crystal boundary at nanometer resolution. No intermediate medium, such as a lipid coating or shroud, could be detected surrounding the crystals. The present study describes a unique application of CSTET in the study of malaria. The findings can be extended to evaluate new drug candidates affecting hemozoin crystal growth.

## Introduction

Malaria remains one of the most serious and life-threatening infectious diseases in the present times. The global burden of the disease estimated in 2020 by the WHO was 241 million cases, with a death toll of 896,000^1^. The causative agent of malaria is a unicellular protozoan parasite-*Plasmodium*, which is transmitted by the bite of an infected female *Anopheles* spp. mosquito. Among the five parasite species that can infect humans, malaria caused by *P. falciparum* is the most fatal. The high mortality rate is attributed to multiple complications, of which cerebral malaria (CM) is the most severe^2^. Even when CM improves upon treatment with potent antimalarial drugs, survivors can experience debilitating after-effects such as seizures and neurocognitive deficits, especially in pediatric patients^2–4^.

Within the human host, after a brief asymptomatic phase within liver cells, *P. falciparum* parasites infect and thrive within red blood cells (RBC), giving rise to the clinical phase of malaria. The life cycle within the infected RBC. (iRBC) lasts for about 48 hours (intraerythrocytic developmental cycle or IDC)^5^. During this time, the parasite undergoes distinct developmental transformations, i.e., ring, trophozoite, and schizont stages (Fig 1A), throughout which the developing parasite derives most of its nutrition from hemoglobin within a digestive vacuole (DV)^6,7^. Catabolism of hemoglobin results in the assimilation of amino acids^8^ and the release of the iron-containing heme^9,10^. The released heme is toxic to the parasite as it can catalyze the formation of reactive oxygen species causing damage to lipids and proteins^11^. Instead, heme is rapidly detoxified by sequestration to physiologically insoluble hemozoin crystals within the DV^12,13^. Hemozoin is often referred to as the ‘malaria pigment’. Hemozoin crystals remain within the DV until the daughter parasites burst out of iRBC, at which point the crystals spill out into the bloodstream eliciting an immune response in the host ^14^.

**Fig 1:**
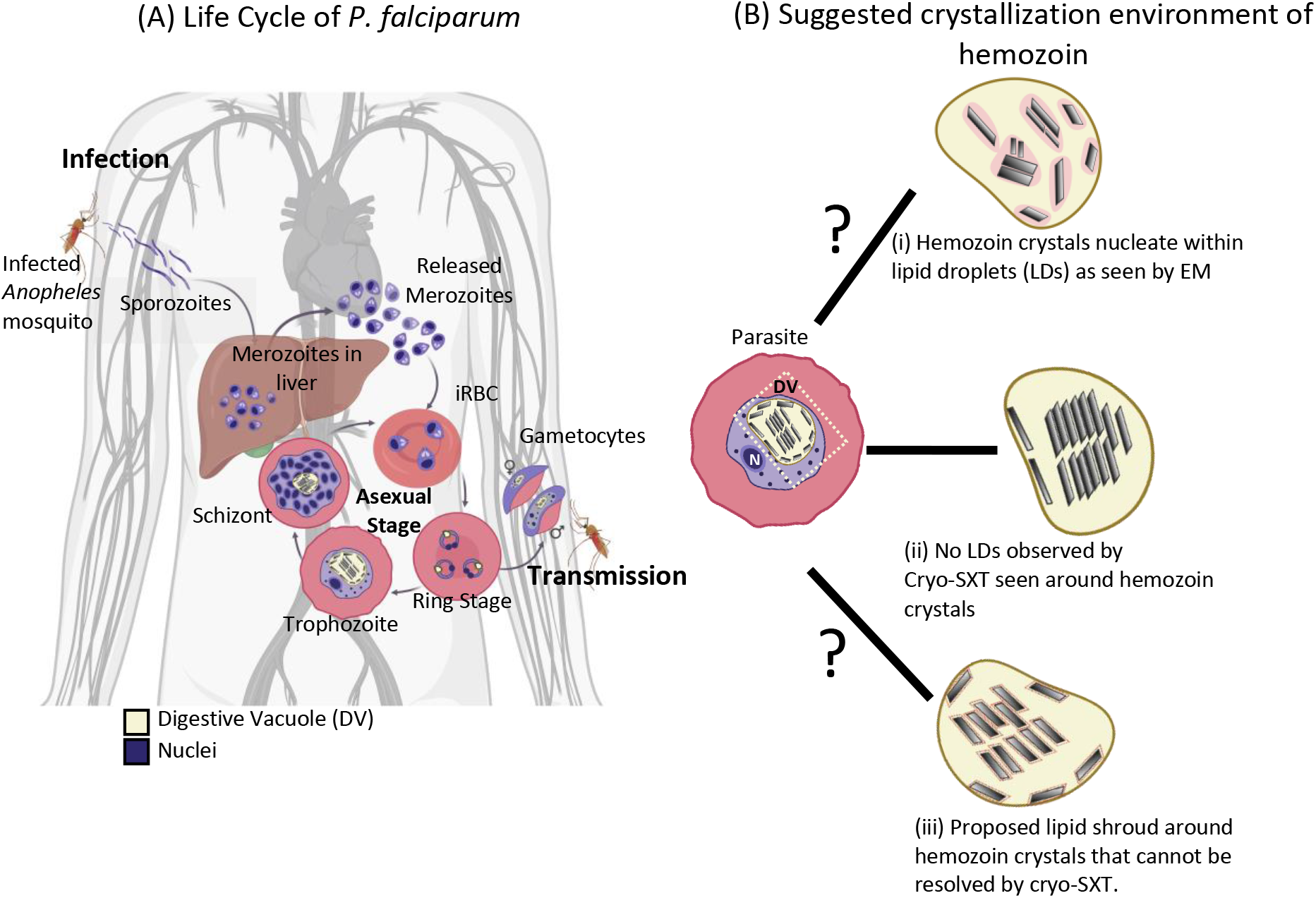
Life-cycle of *Plasmodium falciparum* and previously proposed crystallization environment of hemozoin. The parasite has a complex life-cycle with several morphologically distinct developmental stages. **(A)** After the release of sporozoites from the bite of the female *Anopheles* mosquito, an initial round of amplification occurs in the liver cells to produce infective daughter cells (merozoites). These infect healthy RBCs, initiating clinical malaria. Within the infected RBC (iRBC) undergo repetitive asexual division cycles occurring every 48 hours and characterized by ring, trophozoite, schizont stages (asexual) and some gametocytes (sexual). Hemoglobin is catabolized during this process to produce the ‘malaria pigment’ or hemozoin crystals– a non-toxic by product from heme sequestration with the digestive vacuole (DV) of the parasite. **(B)** Several suggestions have been made about the crystallization environment of hemozoin. (i) Evidence from malachite green staining before EM suggested that the nucleation and growth of hemozoin would likely be in a lipid droplet (LD). This was accepted given the hydrophobic nature of heme. However, most EM data did not reveal this LD, and this was attributed to harsh preparative protocols in conventional EM (ii) Putative LDs were not observed around hemozoin using cryo-SXT. It was later argued that (iii) there may be a ‘lipid shroud’ around the crystal that cannot be resolved by cryo-SXT.

Disruption of the heme detoxification processes is an important target for many existing antimalarial drugs^15–18^. The spread of resistance to these drugs^19–21^ motivates a search for new treatments.^22^ Given that hemozoin is a chemically fixed structure whose production is both indispensable and unique to the parasite, inhibition of heme detoxification processes still remains an essential drug target^17,23^. An improved understanding of the mechanism of hemozoin formation will be crucial in discovering new drug candidates.

The crystallization environment of hemozoin within the DV has been the subject of a long-standing debate in the field (Fig 1B). At first, hemozoin was considered a polymer that resulted from the sequestration of heme within the DV^24,25^. Later studies on hemozoin with X-Ray diffraction and electron paramagnetic resonance (EPR) confirmed the crystalline nature of hemozoin, which comprises regularly repeated dimers of β-hematin with a (Fe^III^-protoporphyrin-IX)_2_ center^26,27^. Given the hydrophobic nature of heme porphyrin, it was proposed that its crystallization, and before that hemozoin ‘polymerization’ occurs within lipid droplets ^28,29^. Moreover, biochemical fractionations of infected red blood cells (iRBCs) identify neutral lipids associated with the hemozoin crystals ^30,31^. Consistently, *in vitro* assays yielding the synthetic analog β-hematin require a fraction of octanol to dissolve the heme in water^32^. Direct observation of a phospholipid layer surrounding synthetic β-hematin was made with cryogenic transmission electron microscopy (cryo-TEM), and it was suggested that this lipid layer aided the crystal formation^33^.

However, similar observations were not made in hemozoin crystals enclosed within the DV of the parasite. The absence of evidence by electron microscopy for a lipid environment surrounding hemozoin in iRBCs was explained by the harsh protocols used to prepare plastic-embedded blocks. In particular, it was proposed that dehydration using an organic solvent might extract the lipid. Upon treatment with the lipid-stabilizing reagent malachite green, a pool of stained material was detected surrounding hemozoin crystals inside the DV, and this was identified as the lipid droplet ^28^. However, subsequent studies of intact parasites by soft X-ray cryo-tomography (SXT) found no evidence of lipid surrounding the hemozoin in intact parasites ^34–36^. Soft X-ray microscopy and tomography are well suited to whole cell investigations of *Plasmodium*, due to the long penetration depth and the quantitative nature of the contrast^37–39^. SXT is doubly suited to detecting lipid bodies: a) physical fixation by vitrification retains both morphology and composition of the specimen, and b) the image contrast is fundamentally based on atomic absorption of low energy X-rays by carbon. It is impossible to hide lipid if present, so the absence of contrast against water is clear evidence of the absence of lipid surrounding the crystals. Crystals were observed to float in the aqueous environment directly and possibly grow from the DV’s inner membrane ^34^. Indeed, it would be challenging to observe heme crystals with sharp facets had they been immersed within a lipid drop (Refer to Fig 4 Kapishnikov *et al*., 2021)^40^. The spatial resolution of SXT is limited to about 25 nm, however, and therefore it was suggested to reconcile the models by the presence of a thin ‘lipid shroud’ surrounding the crystals ^41,42^.

**Fig 2:**
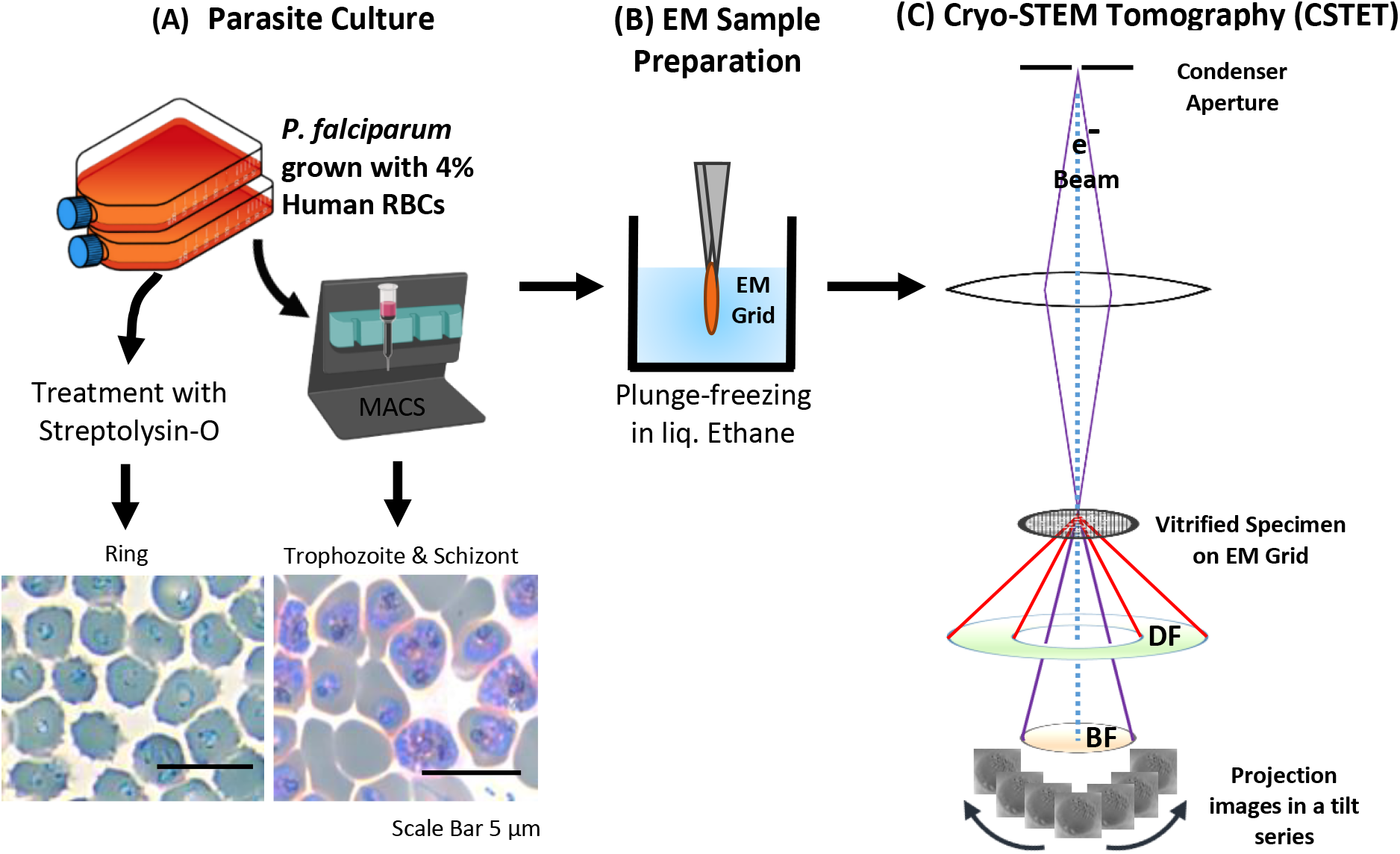
Experimental workflow for using Cryo-STEM tomography (CSTET) to study hemozoin crystals within *P. falciparum* infected RBCs (iRBCs) *P. falciparum* is grown in the lab using standard techniques **(A)** The culture is enriched for iRBCs using a magnetically activated cell-sorting (MACS) column suitable for trophozoites and schizonts and Streptolysin-O (SLO) treatment ring-stage parasites. The enriched iRBCs were smeared and stained with Giemsa to verify the stage before plunging **(B)** Samples were applied to EM grid, blotted and plunge-frozen in liquid ethane **(C)** CSTET makes use of a fine focused electron probe which raster scans the sample, forming an image pixel by pixel. The image is formed by an incoherent process, and the transmitted electrons are detected by detectors kept at different scattering angles (BF= Bright Field; DF=Dark field). For the present study, we have used the information only from the BF detectors. Reconstruction of the tilt-series is done with IMOD.

**Fig 3:**
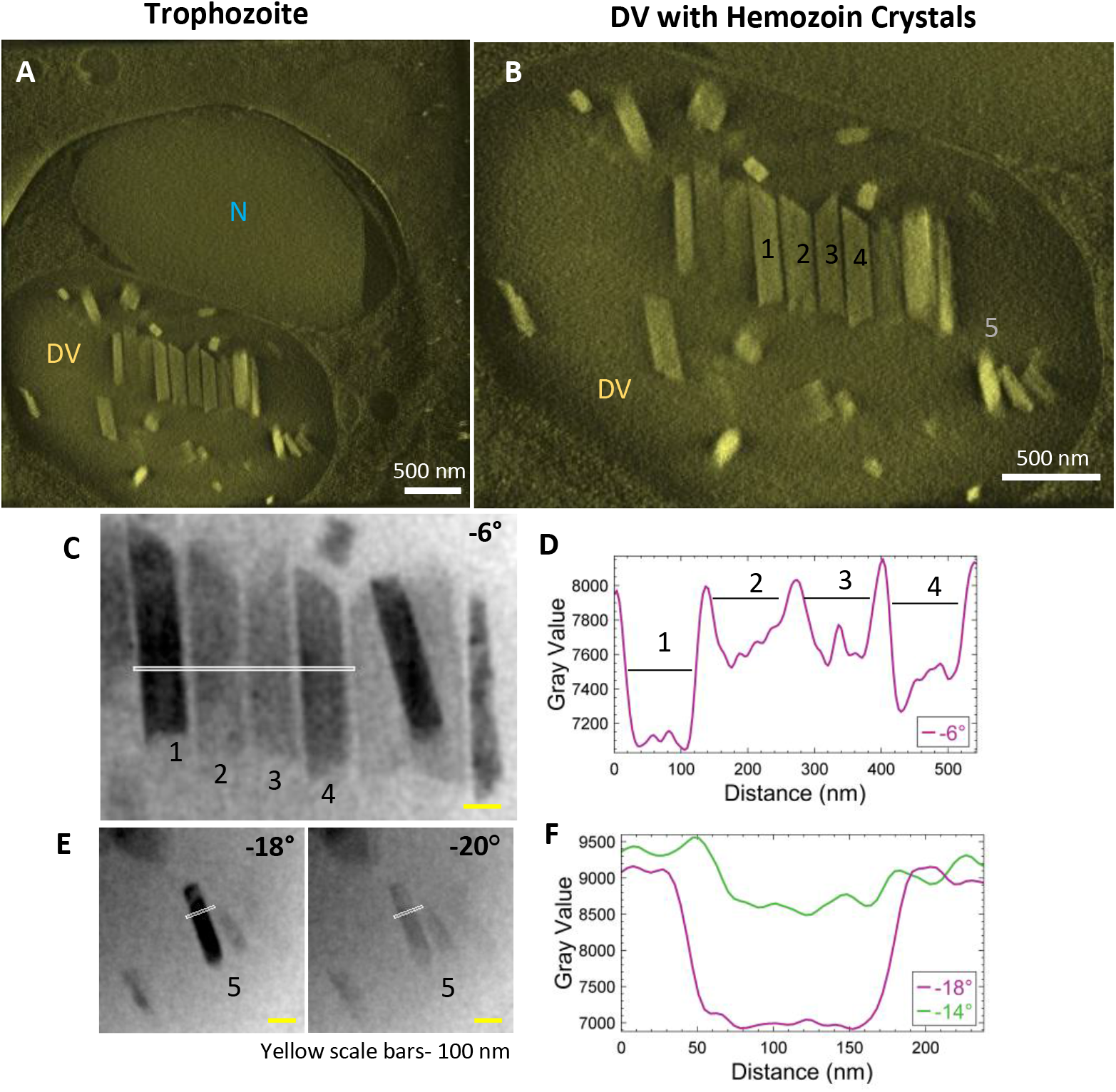
Hemozoin crystals in trophozoite-stage parasites mutually align within the DV and do not possess a lipid shroud around them. Trophozoites are the most metabolically active stage of the parasites, where they actively digest large amounts of hemoglobin **(A)** Thick volume section from the tomogram reconstruction showing the nucleus (N) and the hemozoin crystals within an intact DV. **(B)** The crystals within the DV lie parallel to each other at this stage. The specific crystals that were subsequently analyzed in the aligned tilt-series are labeled from #1-5 and line-intensity profiles were plotted from the white line passing through the crystals **(C)** A series of aligned crystals that go dark and bright (‘blink’) in nearby tilts. Crystals-#1&4 are dark at −6°. **(D)** Line-profile of intensities showing crystal boundaries. **(E)** A smaller crystal (#5) also does not possess a visible lipid shroud. **(F)** Intensity profile across the crystal at adjacent tilts. Note the equal width of the crystal as they go from dark to bright.

**Fig 4:**
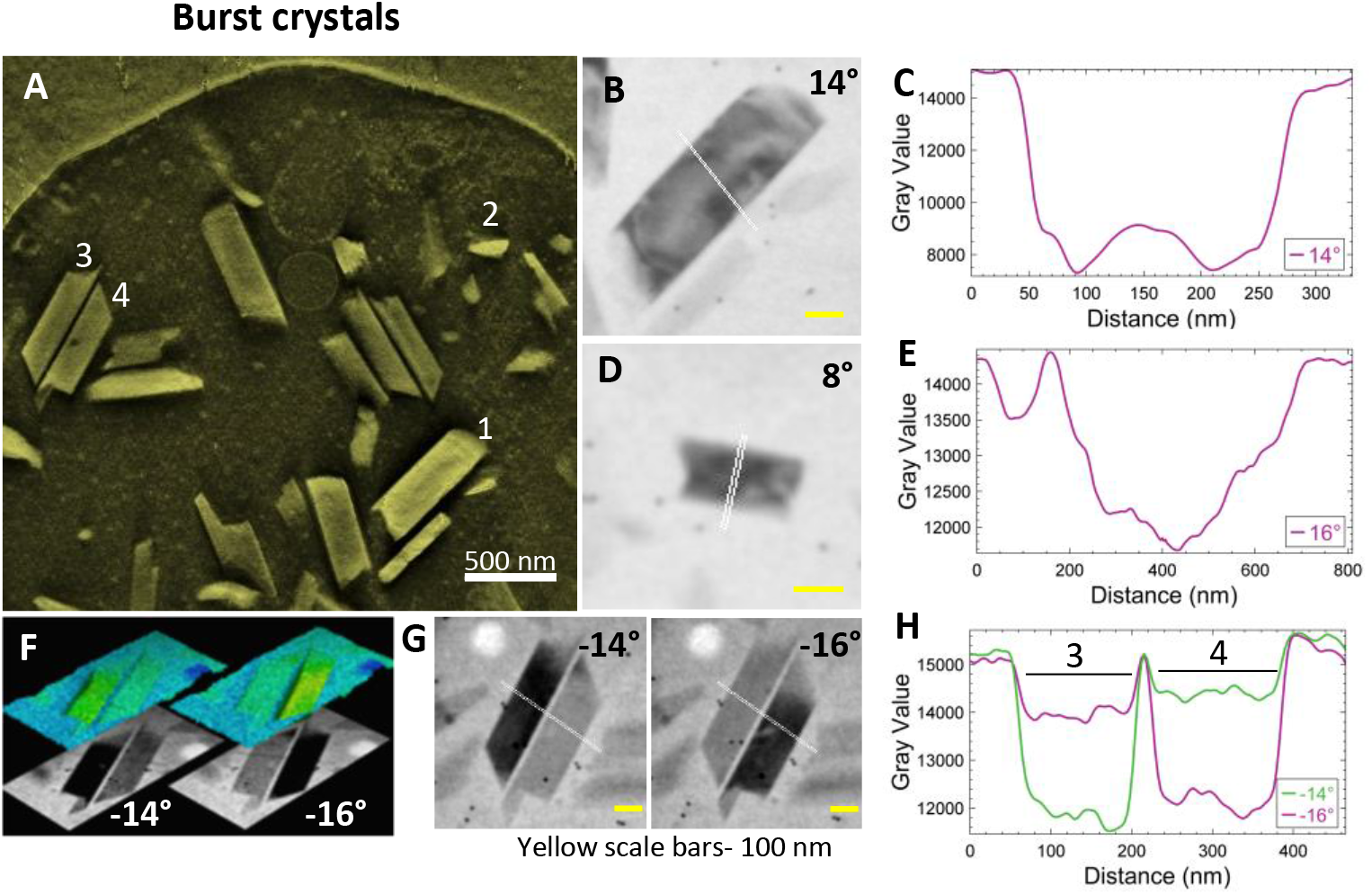
Lipid shroud is absent around cell-free hemozoin crystals. Parasites with their DVs often burst on grids during the blotting process, liberating cell-free crystals that are easier to inspect. **(A)** A thick-volume representation from a reconstruction showing crystals and spilled cellular contents. Crystals that were chosen for analysis are marked #1-4. Projection images from aligned tilt-series are used for analysis. Intensity is plotted along the white-line passing through the crystal. **(B)** Crystal #1 **(C)** Adjoining intensity profile **(D)** Crystal #2 **(E)** Intensity profile. **(F)** Two nearby crystals (#3 & 4) diffract at adjacent tilt angles. Surface topography shows the sharp edges. **(G)** Intensity profiles across the two crystals across the white lines at adjacent tilt angles. The precise fit of the crystal boundaries when light or dark is direct evidence against the presence of an intermediary, non-diffracting layer. **(H)** Intensity profiles across crystals #3-4.

Here we address the nature of the hemozoin-water interface in biogenic hemozoin using an orthogonal cryo-tomography method, Cryogenic Scanning Transmission Electron Tomography (CSTET). This method has been developed in complement to conventional wide-field cryo-electron tomography^43,44^ and is highly sensitive to crystallinity. Based on scanning transmission electron microscopy (STEM), a focused probe rasters across the specimen while transmitted electrons are detected on the opposite side. The contrast reflects the scattering of electrons away from the incident illumination direction. Proper configuration of the STEM modality enables imaging of thick specimens (even 1 μm or more) without the need for plastic embedding or the addition of heavy metal stains. A series of projection images are obtained by tilting the specimen through a range of angles, and three-dimensional reconstructions are generated from the tilt-series.

We used ring and trophozoite stage parasites for CSTET imaging of hemozoin crystals within the DV of intact iRBCs. In the aligned tilt-series, the crystals become suddenly dark at particular tilt angles where Bragg diffraction conditions are met. This sudden change in contrast helps us look for a putative ‘lipid shroud’ surrounding the hemozoin crystals.

## Experimental Section

### 1. Parasite culture and isolation of stages

*Plasmodium falciparum* (NF54) culture was maintained in RPMI supplemented with hypoxanthine and AlbuMAX™ II Lipid-Rich B.S.A. at 4% hematocrit (packed cell volume of RBCs) at 37°C using a gas mixture of 1% oxygen, 5% carbon dioxide in a nitrogenous atmosphere. The parasites were synchronized using sorbitol treatment^45^. The trophozoite stage with well-developed hemozoin was enriched magnetically using MACS^®^ columns (Miltenyi Biotech) ^46^. Ring-stage parasites were selected by Streptolysin–O (SLO) treatment^47^ and given an hour to recover before sample preparation for electron microscopy.

### 2. Preparation of EM samples

Intact iRBCs were applied to standard Quantifoil EM grids having holes of 3.5 μm diameter, blotted with filter paper from behind, and plunged into liquid ethane for vitrification using the Leica EM-GP Plunger (Leica Microsystems). As the hole size is smaller than the iRBC, the cells tend to lodge. Often, they burst, exposing the parasites. Sometimes the parasites also burst, spreading the contents onto the surrounding area. We have used intact as well as burst-out hemozoin for our studies. Focused ion beam (FIB) milling was performed in the Zeiss Crossbeam scanning electron microscope (SEM) to createthin lamellae (300 nm thickness) from iRBCs plunged on 2 μm diameter Quantifoil EM grids.

### 3. CSTET acquisition and analysis

Tomographic series of tilted STEM images were collected using different microscopes -Talos Arctica/ Tecnai T20F (200 kV) or Titan Krios (300 kV) microscopes (Thermo Fisher Scientific). A focused-probe beam of 1.2 mrad semi-convergence angle (~2 nm diameter) was used, over a typical tilt range from −60 to 60 degrees with an interval of 2-degree tilt, with a frame size of 2048 x 2048 pixels. The same imaging parameters were maintained for CSTET while using FIB milled lamellae as well. The series were aligned using IMOD software and reconstructed by weighted back-projection and SIRT-like filter with 30 iterations^48^. It was visualized using UCSF Chimera ^49^ and analyzed using FIJI ^50^.

## Results and Discussion

### 1. Trophozoite-stage hemozoin crystals align together within the DV and are not surrounded by an observable lipid shroud

Synchronized *P. falciparum* cultures were smeared and stained with Giemsa to verify the stage before subjecting to MACS for enrichment. Tilt-series reconstruction from intact, plunge-frozen parasites showed DVs enclosing hemozoin crystals. Fig 3 shows an example of a trophozoite stage parasite with a single large nucleus (Fig 3A; N) with parallel, aligned crystals (Fig 3B; Figs S1 A&B; Movie SV2); some isolated ones dispersed within the DV are also observed. It was suggested that this aligned arrangement of crystals indicates their templated nucleation at a common surface ^51^. The tilt-series could be used to study the crystals and their boundaries directly, even prior to tomographic reconstruction. Bragg reflections cause crystals to appear dark at certain tilt-angles where the scattering away from the bright-field detector is strong (Fig 3C; Crystals #1 and 4 at −6°; Movie SV1). The line-profile for intensities highlights the boundaries of the crystals (Fig 3D). Often, crystals that go dark at one tilt reappear as bright at the next angle (Fig E; Crystal #5 at −18° and −20°, respectively). Note the identical widths in bright and dark states (Fig F). Had there been a surrounding lipid shroud, each crystal should have appeared narrower when dark (diffracting) than when light, as the lipid medium would not meet Bragg conditions (Fig 3D&F).

Additionally, a variety of crystal sizes was observed at this stage, signifying that each crystal was at a different phase of growth. This is expected as the parasites are at their metabolic peak during the trophozoite stage. They continuously catabolize hemoglobin and other host factors for obtaining nutrition^52,53^. It is also interesting to note the asymmetric ends of the crystals, with one end appearing slanted and sharp, and the other broken or ragged. Similar asymmetric shapes have been observed previously in biogenic hemozoin, without comment^54,55^ or with a suggestion that they may represent drug-sensitive growth sites^53^. The fact that the crystals grow with a constant or even decreasing aspect ratio implies that growth occurs in all orientations.

### 2. A visible lipid shroud is not detected in cell-free crystals

Cell-free samples of crystals liberated from burst DVs are thinner than intact cells and provide a higher resolution view. While the bursting is evidently destructive to the cell, the time interval from blotting to plunging is at most a few seconds. Whatever hydrophobic lipid layer the crystals within DV might carry with them would have to be retained upon release into the surrounding aqueous medium. Fig 4A shows a thick volume slice from a reconstructed tomogram containing crystals from a burst cell. A few crystals (#1-4) were selected for closer inspection. Projection images from the aligned tilt-series (Figs 4 B and D) show crystals #1 and 2 blinking dark at 16° and 8°, respectively. The line profiles show the boundaries of the individual crystals (Figs 4 C&E).

Crystals #3 and 4 are a pair of antiparallel hemozoin crystals that are otherwise similar in orientation. They blink conveniently at adjacent stage tilts. These images are compared in Fig 4F as 2D gray level and false-color surface topography. By morphology, the c axis (long-axis)^56^ lies in the plane of view. Bragg diffraction arises when the zonal axis of the crystal is perpendicular to the illumination, i.e., perfectly vertical. Figs 4 G&H shows that the crystals blink at adjacent tilts and that the intensity profiles across the sharp crystal boundaries are clear in Figs 4 E&I. Similar to observations in intact cells, the crystal boundaries do not hint at the presence of a lipid layer.

### 3. Hemozoin crystals are closely packed within the DVs of schizont-stage parasites

Schizonts are characterized by multiple daughter nuclei embedded within a common cytoplasm. Fig 5A shows developing daughter nuclei with an intact DV magnified in Fig 5B (Movie SV3). Large crystals within the DV are closely packed at this stage. As mentioned above, many crystals appear to have a ragged end on one side and a smoother slanted end opposite (Fig 5B; grey arrows). An antiparallel pair of otherwise similar orientations blink conveniently at adjacent stage tilts (Fig 5C), and their intensity profiles are compared (Fig 5D; Movie SV4). The orientation of the crystals is very similar to that observed previously in Fig 4F. The boundaries are clear, and there is no indication of a lipid shroud around them.

**Fig 5:**
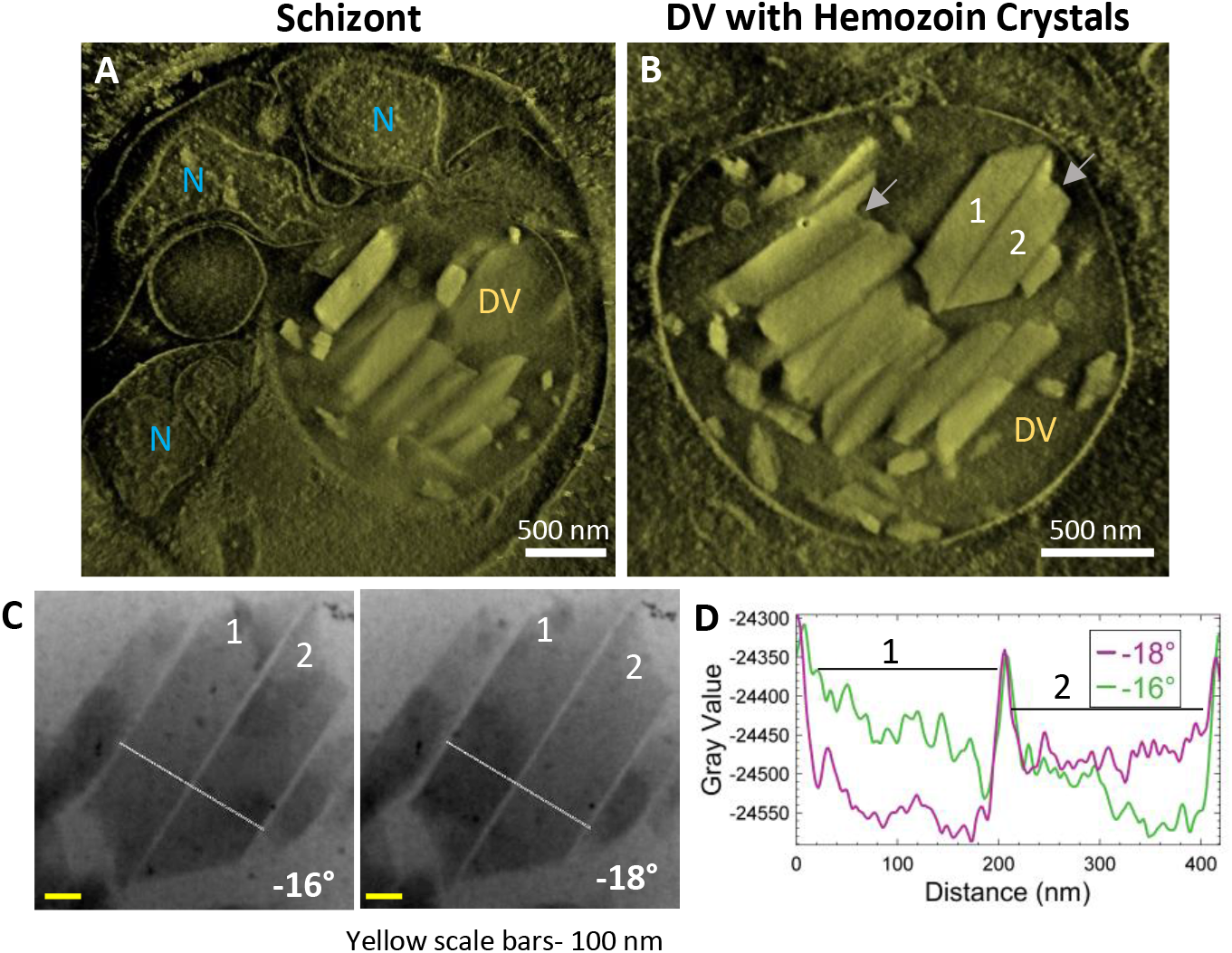
Schizont stage DVs have closely-packed hemozoin crystals within the DV with no visible lipid shroud. Schizonts have multiple daughter nuclei at various stages of development. **(A)** A thick-volume representation from a reconstruction showing crystals within a DV and multiple nuclei (N) **(B)** The DV with many large hemozoin crystals of which a pair (#1 & 2) are analyzed. Note the ragged-end of some of these crystals shown with gray arrows **(C)** Both crystals (#1 & 2) blink dark and bright at adjacent tilts. Intensity is plotted from the white-line passing through the crystal in **(D)** Plotted intensities shows the boundary of the crystal and how they alternately blink at adjacent tilts. The overlapping large central peak is the interface between the two closely-lying crystals.

### 4. Ring-stage parasites have multiple proto-DVs with hemozoin crystals lining the inner membrane

Streptolysin-O (SLO) treated cultures with the enriched ring-stage parasites were plunge-frozen and imaged by CSTET. A thick volume section from the reconstruction in Fig 6A shows the parasite with a single nucleus and multiple small precursors of the DV (proto-DVs) (Fig 6A; Fig S2 A). The hemozoin crystals at this stage are small, and Bragg diffraction is not as apparent as in the later stages. However, adjacent angles from the aligned tilt-series do show flickering hemozoin crystals (Movie SV5). The crystals have a notably higher aspect ratio and are almost exclusively localized to the inner surface of the proto-DV Fig 6B (Movie SV6). This is consistent with previous reports that the initial nucleation occurs on the DV inner membrane and likely involves a lipid medium in some way^34^. The crystal boundaries do not appear as sharp as in the later stages (Fig 6C and D). However, we should recall that the crystal width dimensions are on the order of ten unit cells^56^, and they are imaged in projection through the entire cell thickness (in this example, the cell was 1.12 μm thick).

**Fig 6:**
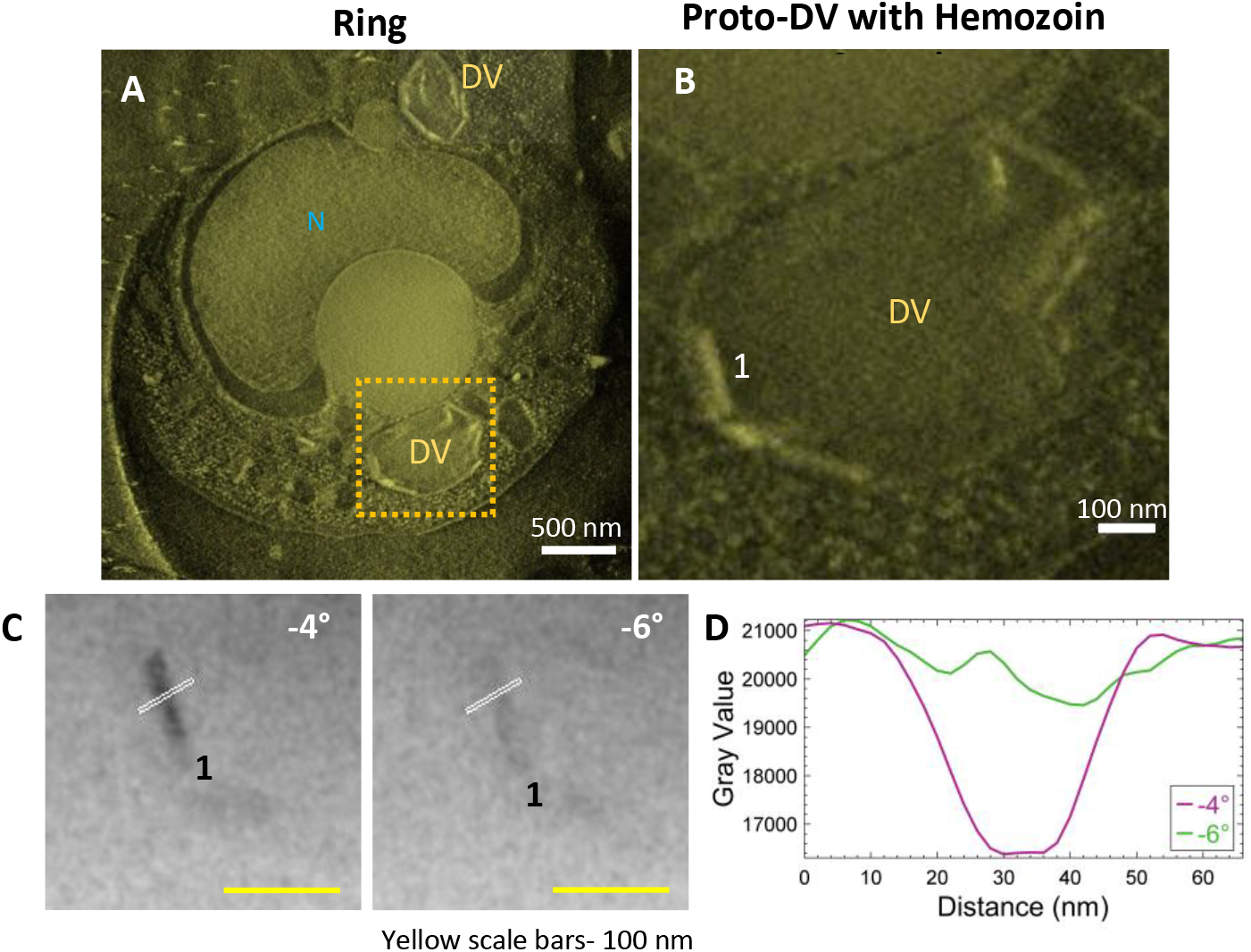
Ring-stage parasites have multiple proto-DVs enclosing small-sized crystals oriented along its inner-membrane. Parasites were treated with SLO for enriching the ring-stage parasites **(A)** A thick-volume representation from a reconstruction showing the nucleus and multiple vacuoles containing hemozoin crystals. **(B)** Small-sized hemozoin crystals lining the inner-membrane of the precursor of the DV (proto-DV). The analyzed crystal is labeled as #1 **(C)** Crystal #1 blinks dark and bright at adjacent tilts. Intensity is plotted from the white-line passing through the crystal in **(D)** Plotted intensities shows the boundary of the crystal.

### 5. The parasite’s endogenous lipid membrane does not show differential contrast across changing angles of the tilt-series

Intact iRBCs can sometimes be too thick to analyze membrane contrast in the raw tilt-series datasets. Reducing the specimen thickness of the vitrified specimen by employing cryo-FIB milling prior to electron tomography can be a potential solution. In the present example (Fig 7), a lamella was milled in the DV of a trophozoite cell to a thickness of 300 nm prior to CSTET. Fig 7A is a thick volume representation of the reconstructed tomogram (Movie SV7) showing the DV enclosing the hemozoin crystals of varied sizes is visible. An example of an endogenous lipid membrane is the DV membrane itself, and we chose two crystals adjacent to it (#1 and 2; Fig 7A). Consistent with data from whole cells, crystal #1 blinks bright and dark between adjacent tilts (Fig 7B). The line profile was limited to the immediate surroundings and showed the expected differences in the intensity (Fig 7C) as observed in previous datasets. Fig 7D shows crystal #2 in a minimum-intensity projection of a thick volume from the reconstruction, together with the neighboring DV membrane (indicated by a red arrow). Intensity line profiles of the projection images were extended to include this membrane as well (Fig 7E). The contrast changes were analyzed across a more extensive range of tilt-angles (−4° to 4°; Fig 7E). While domains of crystal #2 blink across the progressive tilt angles from the aligned tilt-series (Movie SV8), the contrast of the DV membrane remains constant, as indicated by the overlapping troughs in the intensity plots (Fig 7F; red arrow). This validates our assertion that had there been a lipid membrane enclosing hemozoin crystals, it would appear as a feature in the contrast that is insensitive to the Bragg diffraction.

**Fig 7:**
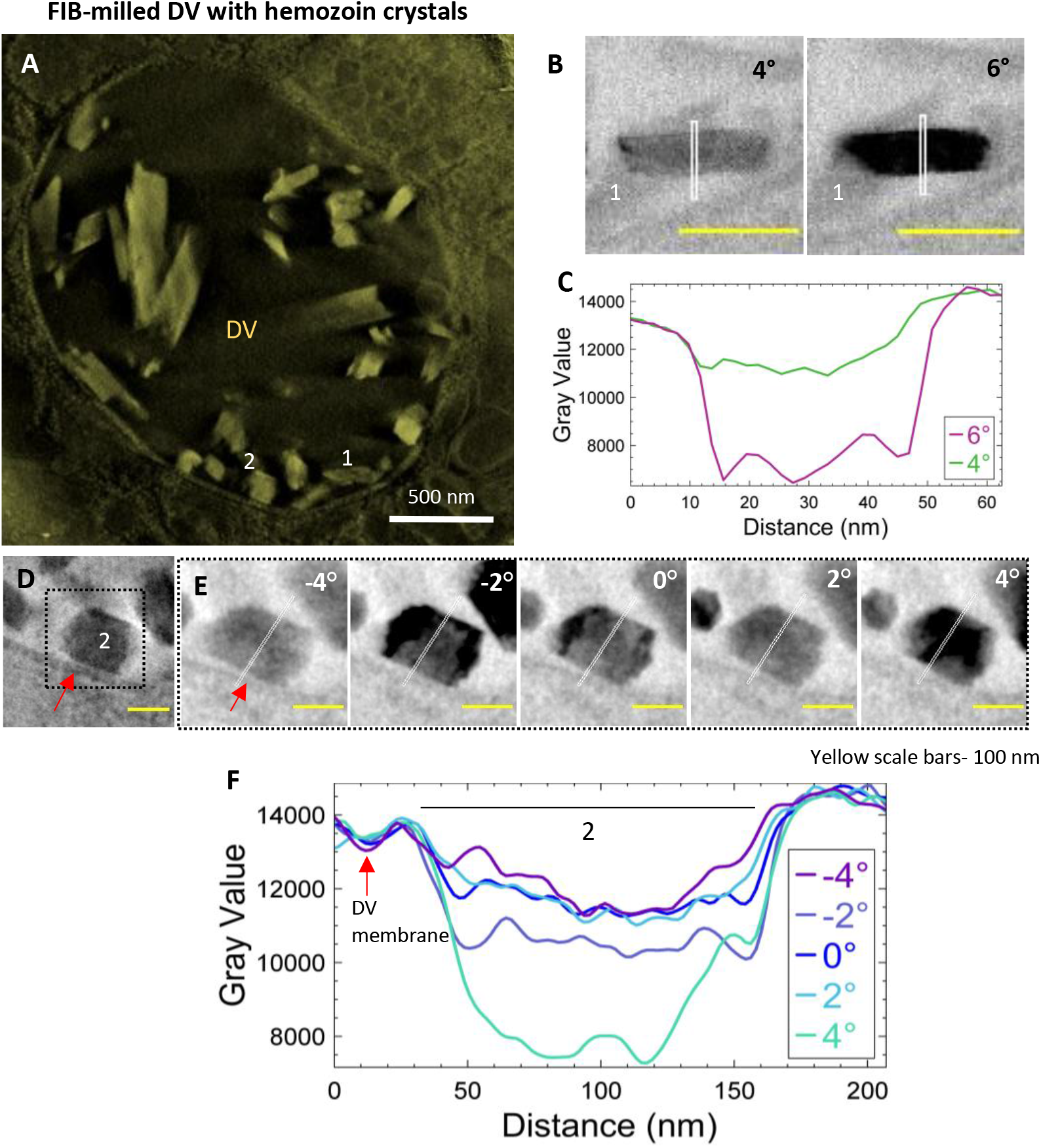
The parasite’s endogenous lipid membrane do not show differential diffraction across changing tilt angles. Parasites on EM grids were cryo-FIB milled to a thickness of 300 nm before CSTET. **(A)** A thick-volume representation from a reconstruction showing the DV enclosing the hemozoin crystals within. #1 and 2 indicate the crystals used for analysis **(B)** Crystal #1 blinks dark and bright at adjacent tilts. Intensity is plotted from the white-line passing through the crystal **(C)** The intensity profile indicates the boundary of the crystal with no detectable lipid shroud. **(D)** Crystal #2 is in the inset indicated by the dotted line with the red arrow pointing to the DV’s membrane (an endogenous lipid membrane) **(E)** Domains of crystal #2 blink across a tilt range from −4° to 4°. The white line passing across the centre of the crystal is extended to include the membrane of the DV while analyzing intensity profiles along these lines. **(F)** Intensity profiles from the tilt angles show the changes, in contrast, observed in crystal #2. In comparison, the intensity of the DV membrane (red arrow) remains more consistent across the same tilts.

## Conclusion

We have systematically studied *P. falciparum* hemozoin crystals *in situ* across the various developmental stages during the asexual phase in iRBCs. During the early phase of infection (ring-stage), the parasite has multiple vacuoles within which the small-sized growing crystals line the inner membrane. It is known that these proto-DVs would fuse to form one large DV in the trophozoite stages^57^. In the trophozoite stage, the crystals may appear aligned side by side within the DV. Additionally, we have analyzed crystals from cells that burst upon blotting of the grid, as well as crystals exposed in cryo-sections formed as FIB-milled lamellae from intact cells. Diffraction contrast reveals the extent of the crystals, which turn suddenly dark when Bragg conditions are met during the tilt series. Had there been an enveloping lipid shroud, the darkened crystal would have to have been enclosed within a wider envelope, similar to observation of the solar corona during an eclipse. From 15 tomographic datasets that we have analyzed in detail, we have never observed a contrast halo surrounding the crystals that might be interpreted as a lipid shroud or any lipid sub-phase in the DV. Antimalarial drugs can thus access the crystal surface directly from the aqueous medium.

## Acknowledgements

DM acknowledges the kind assistance of members of the NRR lab, particularly Anna Rivkin, Yifat Ofir-Birin and Paula Abou Karam, for assistance with parasite culture and handling. Panels of Figs 1, 2 and content entry graphics were created using BioRender.com. This work was supported in part by research grants from the Minerva Research Foundation, the Israel Science Foundation (grant no. 1696/18), and the Estate of David Levinson. Participation of RD was supported by Israel Science Foundation grant no. 1523/18, the Ministry of Science and Technology grant no. 103240 and by the United States-Israel Binational Science Foundation (BSF) grant no. 2019236. RD is also supported by the Dr. Louis M. Leland and Ruth M. Leland Chair in Infectious Diseases. ME is the Head of the Irving and Cherna Moskowitz Center for Nano and Bionano Imaging and incumbent of the Sam and Ayala Zacks Professorial Chair in Chemistry.

## Author Contributions

DM prepared samples and acquired EM data. KR optimized Cryo-FIB lamella production from iRBCs. DM, LL, and ME analyzed data. NR-R and RD provided essential materials and support. DM and ME wrote the paper.

## References

1. World Malaria Report 2021, Geneva: World Health Organization. World Malaria Report 2021. (2021).

2. Schiess, N. et al. Pathophysiology and neurologic sequelae of cerebral malaria. Malar. J. 19, 266 (2020).

3. Mita-Mendoza, N. K. et al. Dimethyl fumarate reduces TNF and Plasmodium falciparum induced brain endothelium activation in vitro. Malar. J. 19, 376 (2020).

4. Boivin, M. J. et al. Cognitive impairment after cerebral malaria in children: a prospective study. Pediatrics 119, e360–6 (2007).

5. Cowman, A. F., Berry, D. & Baum, J. The cellular and molecular basis for malaria parasite invasion of the human red blood cell. J. Cell Biol. 198, 961 LP–971 (2012).

6. Goldberg, D. E., Slater, A. F., Cerami, A. & Henderson, G. B. Hemoglobin degradation in the malaria parasite Plasmodium falciparum: an ordered process in a unique organelle. Proc. Natl. Acad. Sci. U. S. A. 87, 2931–2935 (1990).

7. Bannister, L. H., Hopkins, J. M., Fowler, R. E., Krishna, S. & Mitchell, G. H. A brief illustrated guide to the ultrastructure of Plasmodium falciparum asexual blood stages. Parasitol. Today 16, 427–433 (2000).

8. Bonilla, J. A. et al. Effects on growth, hemoglobin metabolism and paralogous gene expression resulting from disruption of genes encoding the digestive vacuole plasmepsins of Plasmodium falciparum. Int. J. Parasitol. 37, 317–327 (2007).

9. Chou, A. C. & Fitch, C. D. Mechanism of hemolysis induced by ferriprotoporphyrin IX. J. Clin. Invest. 68, 672–677 (1981).

10. Vincent, S. H. Oxidative effects of heme and porphyrins on proteins and lipids. Semin. Hematol. 26, 105–113 (1989).

11. Kumar, S. & Bandyopadhyay, U. Free heme toxicity and its detoxification systems in human. Toxicol. Lett. 157, 175–188 (2005).

12. Jani, D. et al. HDP - A novel heme detoxification protein from the malaria parasite. PLoS Pathog. 4, (2008).

13. Weissbuch, I. & Leiserowitz, L. Interplay Between Malaria, Crystalline Hemozoin Formation, and Antimalarial Drug Action and Design. (1991) doi:10.1021/cr078274t.

14. Olivier, M., Van Den Ham, K., Shio, M. T., Kassa, F. A. & Fougeray, S. Malarial pigment hemozoin and the innate inflammatory response. Front. Immunol. 5, 25 (2014).

15. Olafson, K. N., Ketchum, M. A., Rimer, J. D. & Vekilov, P. G. Mechanisms of hematin crystallization and inhibition by the antimalarial drug chloroquine. 112, (2015).

16. Pandey, A. V., Tekwani, B. L., Singh, R. L. & Chauhan, V. S. Artemisinin, an endoperoxide antimalarial, disrupts the hemoglobin catabolism and heme detoxification systems in malarial parasite. J. Biol. Chem. 274, 19383–19388 (1999).

17. Fong, K. Y. & Wright, D. W. Hemozoin and antimalarial drug discovery. Future Med. Chem. 5, 1437–1450 (2013).

18. Solomonov, I. et al. Crystal nucleation, growth, and morphology of the synthetic malaria pigment β-hematin and the effect thereon by quinoline additives: The malaria pig-ment as a target of various antimalarial drugs [J. am. Chem. Soc. 2007, 129, 2615-2627]. J. Am. Chem. Soc. 129, 5779–5779 (2007).

19. Martin, R. E. et al. Chloroquine transport via the malaria parasite’s chloroquine resistance transporter. Science 325, 1680–1682 (2009).

20. Raj, D. K. et al. Disruption of a Plasmodium falciparum multidrug resistance-associated protein (PfMRP) alters its fitness and transport of antimalarial drugs and glutathione. J. Biol. Chem. 284, 7687–7696 (2009).

21. Suresh, N. & Haldar, K. Mechanisms of artemisinin resistance in Plasmodium falciparum malaria. Curr. Opin. Pharmacol. 42, 46–54 (2018).

22. Yadav, K. et al. Repurposing of existing therapeutics to combat drug-resistant malaria. Biomed. Pharmacother. 136, 111275 (2021).

23. de Villiers, K. A. & Egan, T. J. Heme Detoxification in the Malaria Parasite: A Target for Antimalarial Drug Development. Acc. Chem. Res. 54, 2649–2659 (2021).

24. Slater, A. F. et al. An iron-carboxylate bond links the heme units of malaria pigment. Proc. Natl. Acad. Sci. U. S. A. 88, 325–329 (1991).

25. Sullivan, D. J., Jr, Matile, H., Ridley, R. G. & Goldberg, D. E. A common mechanism for blockade of heme polymerization by antimalarial quinolines. J. Biol. Chem. 273, 31103–31107 (1998).

26. Pagola, S., Stephens, P. W., Bohle, D. S., Kosar, a. D. & Madsen, S. K. The structure of malaria pigment beta-haematin. Nature 404, 307–310 (2000).

27. Sienkiewicz, A. et al. Multi-frequency high-field EPR study of iron centers in malarial pigments. J. Am. Chem. Soc. 128, 4534–4535 (2006).

28. Pisciotta, J. M. & Sullivan, D. Hemozoin: oil versus water. Parasitol. Int. 57, 89–96 (2008).

29. Fitch, C. D., Cai, G.-Z., Chen, Y.-F. & Shoemaker, J. D. Involvement of lipids in ferriprotoporphyrin IX polymerization in malaria. Biochim. Biophys. Acta Mol. Basis Dis. 1454, 31–37 (1999).

30. Kuter, D. et al. Insights into the initial stages of lipid-mediated haemozoin nucleation. CrystEngComm 18, 5177–5187 (2016).

31. Pisciotta, J. M. et al. The role of neutral lipid nanospheres in Plasmodium falciparum haem crystallization. 204, 197–204 (2007).

32. Ketchum, M. A., Olafson, K. N., Petrova, E. V., Rimer, J. D. & Vekilov, P. G. Hematin crystallization from aqueous and organic solvents. The Journal of Chemical Physics 139, 121911 (2013).

33. Huy, N. T. et al. Phospholipid Membrane-Mediated Hemozoin Formation: The Effects of Physical Properties and Evidence of Membrane Surrounding Hemozoin. PLOS ONE 8, e70025 (2013).

34. Kapishnikov, S. et al. Oriented nucleation of hemozoin at the digestive vacuole membrane in Plasmodium falciparum. PNAS 109, 11188–11193 (2012).

35. Kapishnikov, S. et al. Biochemistry of malaria parasite infected red blood cells by X-ray microscopy. Scientific Reports 7, 802 (2017).

36. Kapishnikov, S. et al. Unraveling heme detoxification in the malaria parasite by in situ correlative X-ray fluorescence microscopy and soft X-ray tomography. Sci. Rep. 7, 1–12 (2017).

37. Hale, V. L. et al. Parasitophorous vacuole poration precedes its rupture and rapid host erythrocyte cytoskeleton collapse in Plasmodium falciparum egress. Proc. Natl. Acad. Sci. U. S. A. 114, 3439–3444 (2017).

38. Hanssen, E. et al. Soft X-ray microscopy analysis of cell volume and hemoglobin content in erythrocytes infected with asexual and sexual stages of Plasmodium falciparum. J. Struct. Biol. 177, 224–232 (2012).

39. Magowan, C. et al. Intracellular structures of normal and aberrant Plasmodium falciparum malaria parasites imaged by soft x-ray microscopy. Proc. Natl. Acad. Sci. U. S. A. 94, 6222–6227 (1997).

40. Kapishnikov, S., Hempelmann, E., Elbaum, M., Als-Nielsen, J. & Leiserowitz, L. Malaria Pigment Crystals: The Achilles’ Heel of the Malaria Parasite. ChemMedChem 16, 1515–1532 (2021).

41. Olafson, K. N., Nguyen, T. Q. & Rimer, J. D. Antimalarials inhibit hematin crystallization by unique drug–surface site interactions. Proceedings of the (2017).

42. Vekilov, P. G., Rimer, J. D., Olafson, K. N. & Ketchum, M. A. Lipid or aqueous medium for hematin crystallization? CrystEngComm 17, 7790–7800 (2015).

43. Elbaum, M. Expanding horizons of cryo-tomography to larger volumes. Curr. Opin. Microbiol. 43, 155–161 (2018).

44. Wolf, S. G., Houben, L. & Elbaum, M. Cryo-scanning transmission electron tomography of vitrified cells. Nat. Methods 11, 423–428 (2014).

45. Lambros, C. & Vanderberg, J. P. Synchronization of Plasmodium falciparum erythrocytic stages in culture. J. Parasitol. 65, 418–420 (1979).

46. Ribaut, C. et al. Concentration and purification by magnetic separation of the erythrocytic stages of all human Plasmodium species. Malar. J. 7, 1–5 (2008).

47. Jackson, K. E. et al. Selective permeabilization of the host cell membrane of *Plasmodium falciparum* - infected red blood cells with streptolysin O and equinatoxin II. Biochem. J 403, 167–175 (2007).

48. Kremer, J. R., Mastronarde, D. N. & McIntosh, J. R. Computer visualization of three-dimensional image data using IMOD. J. Struct. Biol. 116, 71–76 (1996).

49. Pettersen, E. F. et al. UCSF Chimera - A visualization system for exploratory research and analysis. J. Comput. Chem. 25, 1605–1612 (2004).

50. Schindelin, J. et al. Fiji: an open-source platform for biological-image analysis. Nat. Methods 9, 676–682 (2012).

51. Kapishnikov, S. et al. Aligned hemozoin crystals in curved clusters in malarial red blood cells revealed by nanoprobe X-ray Fe fluorescence and diffraction. Proc. Natl. Acad. Sci. U.S.A. 109, 11184–11187 (2012).

52. Counihan, N. A., Modak, J. K. & de Koning-Ward, T. F. How malaria parasites acquire nutrients from their host. Front. Cell Dev. Biol. 9, 649184 (2021).

53. Wendt, C. et al. High-Resolution Electron Microscopy Analysis of Malaria Hemozoin Crystals Reveals New Aspects of Crystal Growth and Elemental Composition. Cryst. Growth Des. 21, 5521–5533 (2021).

54. Ambele, M. A., Sewell, B. T., Cummings, F. R., Smith, P. J. & Egan, T. J. Synthetic Hemozoin (β-Hematin) Crystals Nucleate at the Surface of Neutral Lipid Droplets that Control Their Sizes. Cryst. Growth Des. 13, (2013).

55. Noland, G. S., Briones, N. & Sullivan, D. J., Jr. The shape and size of hemozoin crystals distinguishes diverse Plasmodium species. Mol. Biochem. Parasitol. 130, 91–99 (2003).

56. Straasø, T. et al. The role of the four stereoisomers of the heme Fe–O cyclic dimer in the crystalline phase behavior of synthetic hemozoin: Relevance to native hemozoin crystallization. Cryst. Growth Des. 11, 3342–3350 (2011).

57. Dluzewski, A. R. et al. Formation of the food vacuole in Plasmodium falciparum: a potential role for the 19 kDa fragment of merozoite surface protein 1 (MSP1(19)). PLoS One 3, e3085 (2008).

