## Supplementary Information for "Diffraction contrast in cryo-scanning transmission electron tomography reveals the boundary of hemozoin crystals *in situ*"

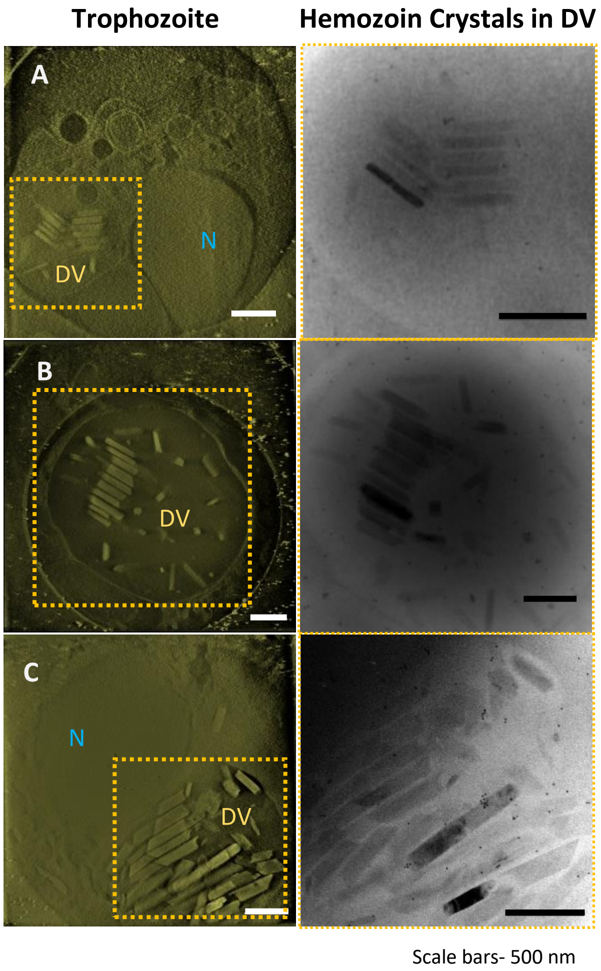


**Fig S1: Hemozoin crystals remain mutually aligned within the DV of trophozoite-stage parasites** The parasites were plunge-frozen and imaged using CSTET. The tilt-series were reconstructed using IMOD and a slice from a thick volume is shown in panel (i) is helpful to identify cellular features like the digestive vacuole (DV; inset shown in panel ii) and nucleus (N). Projection images from the aligned tilt-series shown in panel (ii) were used for observation of Bragg diffraction across specific tilt-angles. **(A)** A trophozoite with mutually aligned hemozoin crystals that go dark and bright in a single slice from the tilt-series crystals are observed **(B)** Another instance of a parasite with mutually aligned hemozoin crystals visibly larger than the previous one. A single tilt with crystals that are dark and bright **(C)** Likely a later trophozoite with a large nucleus and crystals tightly-packed within a DV.

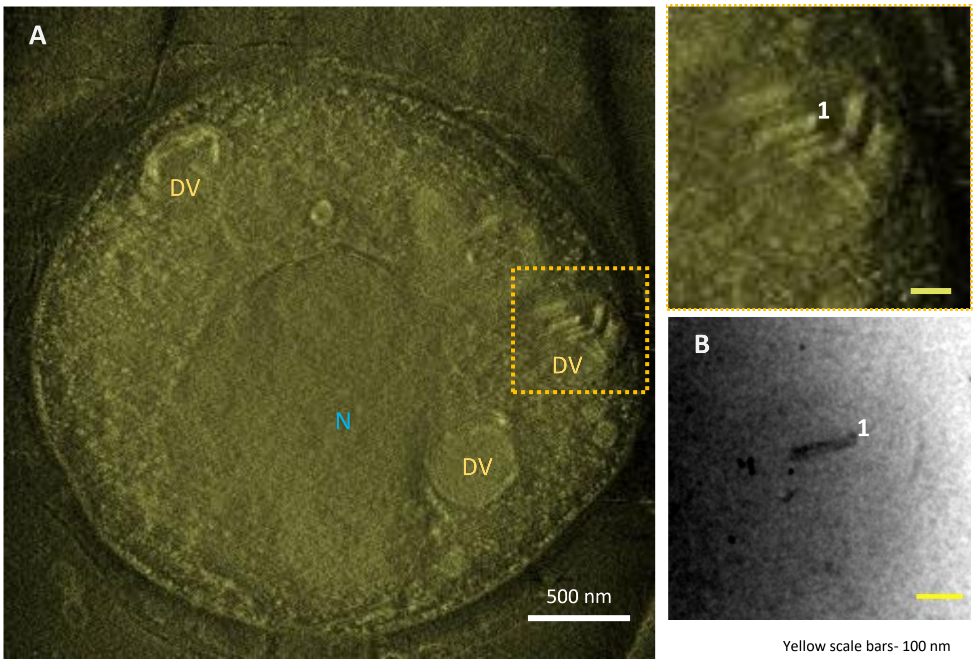


**Fig S2: Ring-stage parasite have multiple proto-DVs enclosing small-sized crystals**
**(A)** A thick-volume slice from a reconstruction showing a single nucleus and multiple spherical proto-DVs. The inset is of a single proto-DV **(B)** A crystal that appears dark in a slice of a tilt-series.

### Supplementary Movies

The aligned tilt-series is normalized for intensities to ensure that the crystals are visible in the higher tilts.

SV1: Trophozoite aligned tilt-series of the DV- G4__BF_ali_Crop.norm.avi

SV2: Trophozoite reconstruction of DV- G4__BF_bin2_rec_Z106-348_GBp75.avi

SV3: Schizont aligned tilt-series of the DV- DM1_tomo1_BF_ali_Bin2.norm.avi

SV4: Schizont reconstruction of DV- DM1_tomo1_BF_bin2_rec_Crop_GBp75.avi

SV5: Ring-stage aligned tilt-series of the DV- DM126_3_Rings_Cell_5_1_ali_Bin2.norm.avi

SV6: Ring-stage reconstruction of DV- DM126_3_Rings_Cell_5_1_rec_bin2_z138_425.avi

SV7: Reconstruction cryo-FIB milled CSTET data of DV- DM48_Tomo_2_BF_rec_bin2_smooth.avi

SV8: Aligned tilt-series from cryo-FIB milled lamella- DM48_Tomo_2_BF_ali_bin2_Z2-61.norm.avi
